# The cell division factor ZapB is required for bile resistance in *Salmonella enterica*

**DOI:** 10.1101/759068

**Authors:** Sara B. Hernández, Rocío Fernández-Fernández, Elena Puerta-Fernández, Verónica Urdaneta, Josep Casadesús

**Author notes:** Department of Molecular Biology, Umeå University, Umeå 90187, Sweden (Sara B. Hernández). Instituto de Recursos Naturales y Agrobiología de Sevilla (IRNAS, CSIC), Avda. Reina Mercedes, 10, 41012 Sevilla, Spain. Department of Infectious Diseases, Yale School of Medicine, The Anylan Center 300 Cedar Street, New Haven 06519, Connecticut, U. S. A. Address correspondence to: Josep Casadesús, Correspondence may be also addressed to: Verónica Urdaneta.

## Abstract

A gene annotated as *yiiU* in the genome of *Salmonella enterica* serovar Typhimurium encodes a protein homologous to *E. coli* ZapB, a non-essential cell division factor involved in Z-ring assembly. ZapB^−^ null mutants of *S. enterica* are bile-sensitive. The ZapB protein is degraded in the presence of sodium deoxycholate (DOC), and degradation appears to involve the Lon protease. The amount of *zapB* mRNA increases in the presence of a sublethal concentration of DOC. This increase is not caused by upregulation of *zapB* transcription but by increased stability of *zapB* mRNA. DOC-induced increase of the *zapB* transcript is suppressed by an *hfq* mutation, suggesting the involvement of a small regulatory RNA. We provide evidence that such sRNA is MicA. Increased stability of *zapB* mRNA in the presence of DOC may counter degradation of bile-damaged ZapB, thus providing sufficient level of functional ZapB protein to permit Z-ring assembly in the presence of bile.

**IMPORTANCE:** Bile salts have bactericidal activity as a consequence of membrane disruption, protein denaturation and DNA damage. However, intestinal bacteria are resistant to bile. Envelope structures such as the lipopolysaccharide and the enterobacterial common antigen act as barriers that reduce intake of bile salts. Remodelling of the outer membrane and the peptidoglycan, activation of efflux pumps, and upregulation of stress responses also contribute to bile resistance. This study adds the cell division factor ZapB (and presumably the Z-ring) to the list of cellular functions involved in bile resistance.

## INTRODUCTION

During infection of the gastrointestinal tract, enteric pathogens must endure harsh conditions such as acidic pH, low oxygen, elevated osmolarity, and nutrient limitation (1, 2). In the small intestine, secretion of bile poses another challenge due to the antibacterial activity of bile salts (3). *Salmonella* serovars that cause systemic and chronic infection also encounter bile in the gall bladder, at concentrations higher and more steady than in the intestine (3, 4). Bile acids/salts act as detergents that disrupt membrane phospholipids, cause misfolding and denaturation of proteins, damage DNA, and interfere with formation of secondary RNA structures (5–8).

Resistance to bile can be studied under laboratory conditions by adding ox bile or individual bile salts to microbiological culture media. In both *E. coli* and *Salmonella*, isolation of bile-sensitive mutants has proven useful to identify loci required for bile resistance, and suppressor analysis has contributed to understand the mechanisms involved (6–8). Bacterial responses to bile have been also investigated by high throughput analysis of gene expression (9, 10). Envelope structures such as the lipopolysaccharide (11–13) and the enterobacterial common antigen (14) provide barriers that reduce intake of bile salts. In addition, bile salts are transported outside the cell by efflux pumps, especially AcrAB-TolC (15–17). Exposure to bile also activates stress responses (10, 18), peptidoglycan remodelling (19) and DNA repair mechanisms (20, 21).

Among the bile-upregulated genes identified by transcriptomic analysis, a *Salmonella enterica* locus annotated as *yiiU* was found (10). Here, we show that *S. enterica* YiiU is a homolog of the *E. coli* ZapB protein, a non-essential cell division factor involved in Z-ring formation and nucleoid segregation (22–25). We also show that *Salmonella* ZapB^−^ mutants are sensitive to sodium deoxycholate (DOC), indicating that ZapB plays a role in bile resistance. The ZapB protein is degraded by the Lon protease in the presence of DOC, probably as a consequence of DOC-induced damage. However, the stability of *zapB* mRNA increases in the presence of DOC. We provide evidence that increased stability may result from the interaction of *zapB* mRNA with MicA, a small regulatory RNA whose synthesis is upregulated by bile.

## RESULTS

### *S. enterica* YiiU is a homolog of the *E. coli* cell division factor ZapB

The *zapB* gene of *E. coli* encodes an 81 amino acid protein that acts as a non-essential cell division factor involved in formation of the Z-ring. The name *zapB* is for ‘Z ring-associated protein B’ (22). Alignment of the amino acid sequences of the ZapB protein of *E. coli* and the predicted protein encoded by the *yiiU* locus of *Salmonella enterica* reveals 91% identity. For this reason, the *S. enterica yiiU* locus will be henceforth named *zapB*.

To detect *S. enterica* ZapB by Western immunoblot analysis, the ZapB protein was tagged with a 3×FLAG epitope. Electrophoretic separation of cell fractions (cytosol, cytoplasmic membrane and outer membrane) and Western analysis of the resolved protein extracts indicated that *Salmonella* ZapB is a cytoplasmic protein like its *E. coli* homolog (Fig. 1A).

**Figure 1.**
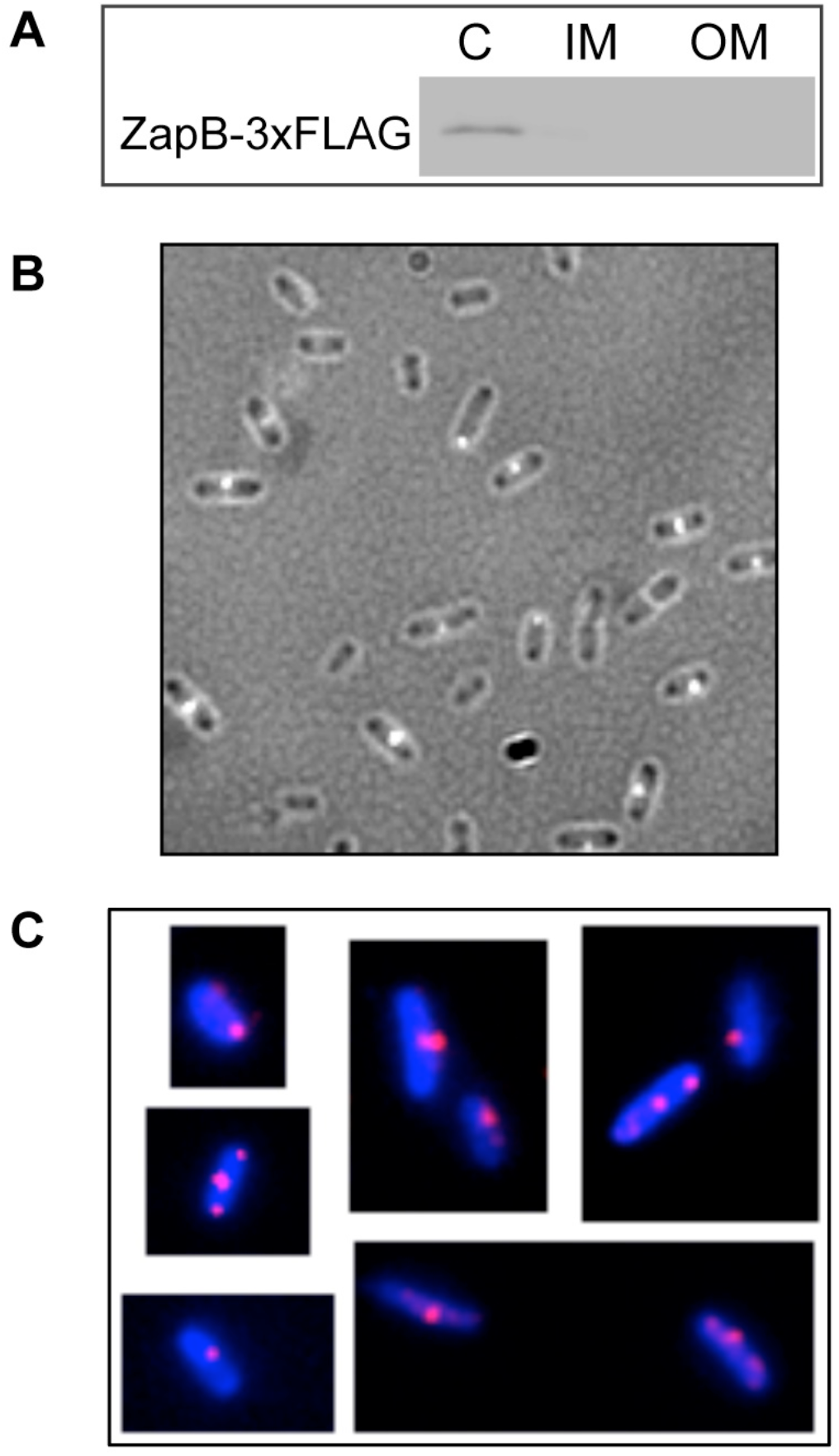
Cellular localization of ZapB. **A.** Distribution of ZapB protein tagged with a 3×FLAG epitope in subcellular fractions of *S. enterica* serovar Typhimurium (strain SV6898). Anti-FLAG western hybridization is shown for three fractions: cytoplasm (C), inner membrane (IM), and outer membrane (OM). The volumes loaded for each fraction were normalized to the same number of bacteria (5×10^8^ CFU). **B.** Combined phase-contrast and fluorescence microscopy of exponentially growing cells of a strain harboring a chromosomal *zapB*::mCherry fusion (SV7571). **C.** Fluorescence images of *zapB*::3×FLAG immunostaining with Hoechst 33342 under the same growth conditions.

To investigate the subcellular localization of *S. enterica* ZapB, a *zapB*-mCherry fusion was constructed on the chromosome. Fluorescence microscopy imaging showed the mCherry signal at the constriction site as previously reported for *E.coli* ZapB (22, 24, 26, 27). In certain cells the mCherry signal was also detected at the poles (Fig. 1B). A caveat in these experiments was that the strain carrying the *zapB*-mCherry construct was sensitive to DOC, suggesting that its intracellular role might be impaired (Table 1). Hence, we also determined the subcellular localization of ZapB in an immunostaining assay with a Cy3 anti-rabbit conjugate antibody using the ZapB-3×FLAG construct described above. This construct was considered *bona fide* wild type because the MIC of DOC of the ZapB-3×FLAG strain (SV6898) was identical to that of the wild type (Table 1). Immunostaining showed ZapB foci at the constriction site, and in certain cells also at the poles (Fig. 1C).

**Table 1.** Minimal concentrations of sodium deoxycholate (DOC)ª

| Strain | Genotype | MIC of DOC (%) |
| --- | --- | --- |
| SL1344 | Wild type | 7 |
| SV6269 | $\Delta zapB$ | 2 |
| SV7264 | SL1344 / pBAD18 | 7 |
| SV7265 | $\Delta zapB$ / pBAD18 | 2 |
| SV7267 | SL1344 / pBAD18::*zapB* | 7 |
| SV7268 | $\Delta zapB$ / pBAD18::*zapB* | 7 |
| SV7432 | $\Delta zapB$ / pBAD18::*zapB-E. coli* | 7 |
| SV6898 | *zapB*::3×FLAG | 7 |
| SV7571 | *zapB*::mCherry | 2 |
| SV7016 | *lon*::Tn10 | 9 |
| SV7022 | $\Delta zapB$ *lon*::Tn10 | 2 |

<sup>a</sup> Medians of >3 experiments

### ZapB is required for bile resistance

Minimal inhibitory concentration (MIC) analysis was performed to determine whether the *S. enterica* ZapB protein is necessary for bile resistance. A null ZapB^−^ mutant (SV6269) was found to be sensitive to DOC (Table 1). The conclusion that ZapB is necessary for bile resistance was confirmed by complementation analysis with plasmid-borne *E. coli* and *S. enterica zapB* genes expressed from the arabinose-inducible promoter of pBAD18 (Table 1).

### ZapB is degraded by the Lon protease in the presence of bile

The stability of the ZapB protein in the presence and in the absence of DOC was analyzed by Western blot analysis using a ZapB version tagged with the 3×FLAG epitope (ZapB::3×FLAG). The amount of ZapB protein in LB + DOC was found to be about half than the amount found in LB (Fig. 2A), suggesting that ZapB might be degraded in the presence of DOC. Protein stability assays were performed to monitor the half-life of the ZapB protein upon addition of DOC. As shown in Fig. 2B (top), ZapB is degraded faster in LB + DOC (half-life around 30 min) than in LB (half-life >90 min).

**Figure 2.**
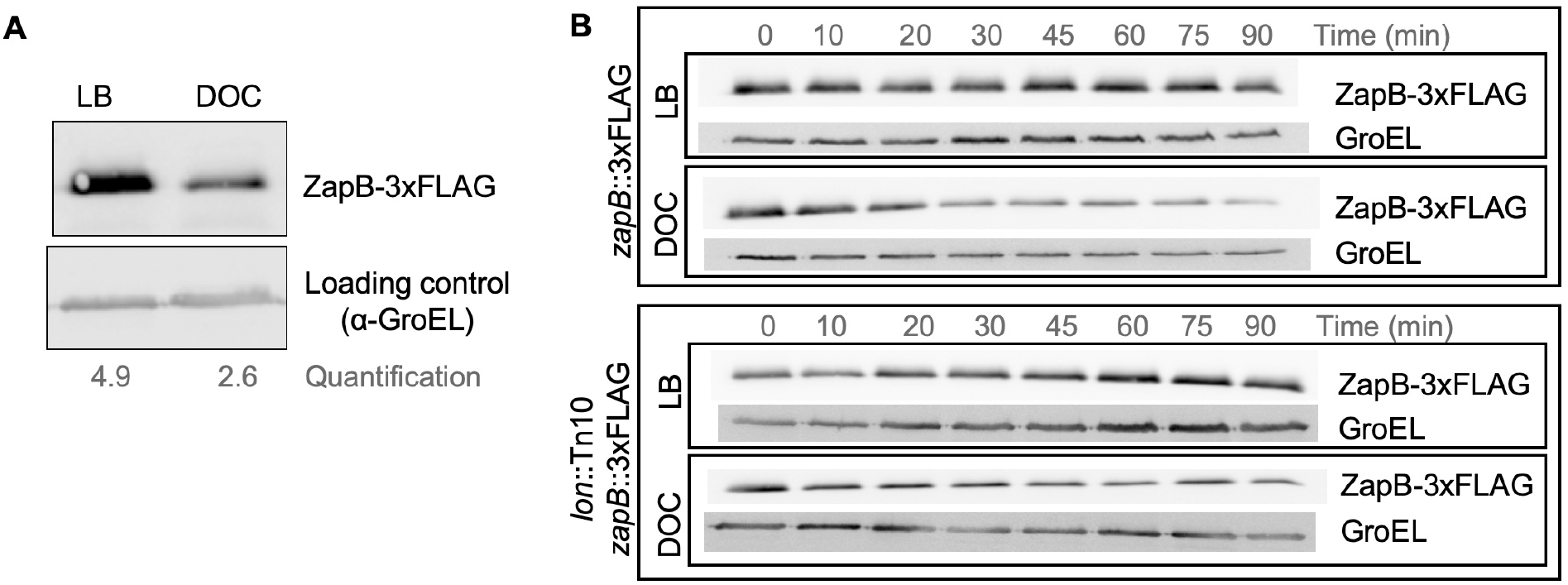
Degradation of ZapB protein by the Lon protease. **A.** Western blot analysis of the levels of ZapB-3×FLAG protein in LB and LB + 5% DOC. **B.** Stability of the ZapB-3×FLAG protein in exponential cultures grown in LB and LB + 5% DOC in wild type and Lon^−^ backgrounds. Aliquots were extracted 10, 20, 30, 45, 60, 75, and 90 min after addition of DOC and chloramphenicol.

Because the Lon protease is involved in the degradation of unfolded and misfolded proteins (28, 29) and bile salts cause misfolding and denaturation of proteins (18, 30), the stability of ZapB protein was monitored in a Lon^−^ background (strain SV7017). In the absence of Lon protease, ZapB was found to be more stable in the presence of DOC (Fig. 2B, bottom). Hence, we tentatively conclude that the Lon protease may degrade ZapB in the presence of DOC.

### Postranscriptional regulation of *zapB*

Transcriptomic analysis had detected upregulation of *zapB* expression in the presence of DOC (10). Upregulation was confirmed by β-galactosidase assays using a translational *zapB::lacZ* fusion (strain SV6270) and by Northern blot analysis (Fig. 3, A and B). To investigate the underlying mechanism, the transcriptional start site of the *zapB* gene was identified by 5’ rapid amplification of 5’ cDNA ends (5’-RACE). Twenty-four out of twenty-nine clones analyzed from two independent 5’-RACE assays identified a 5’-end A residue located 56 nucleotides upstream of the *zapB* translational start codon. *In silico* analysis of the region revealed the occurrence of nucleotide sequences compatible with the consensus sequences of −35 and −10 regions in σ^70^-dependent promoters, and with appropriate spacing (31). The proposed structure of the *zapB* promoter is presented in Suppl. Fig. S1.

**Figure 3.**
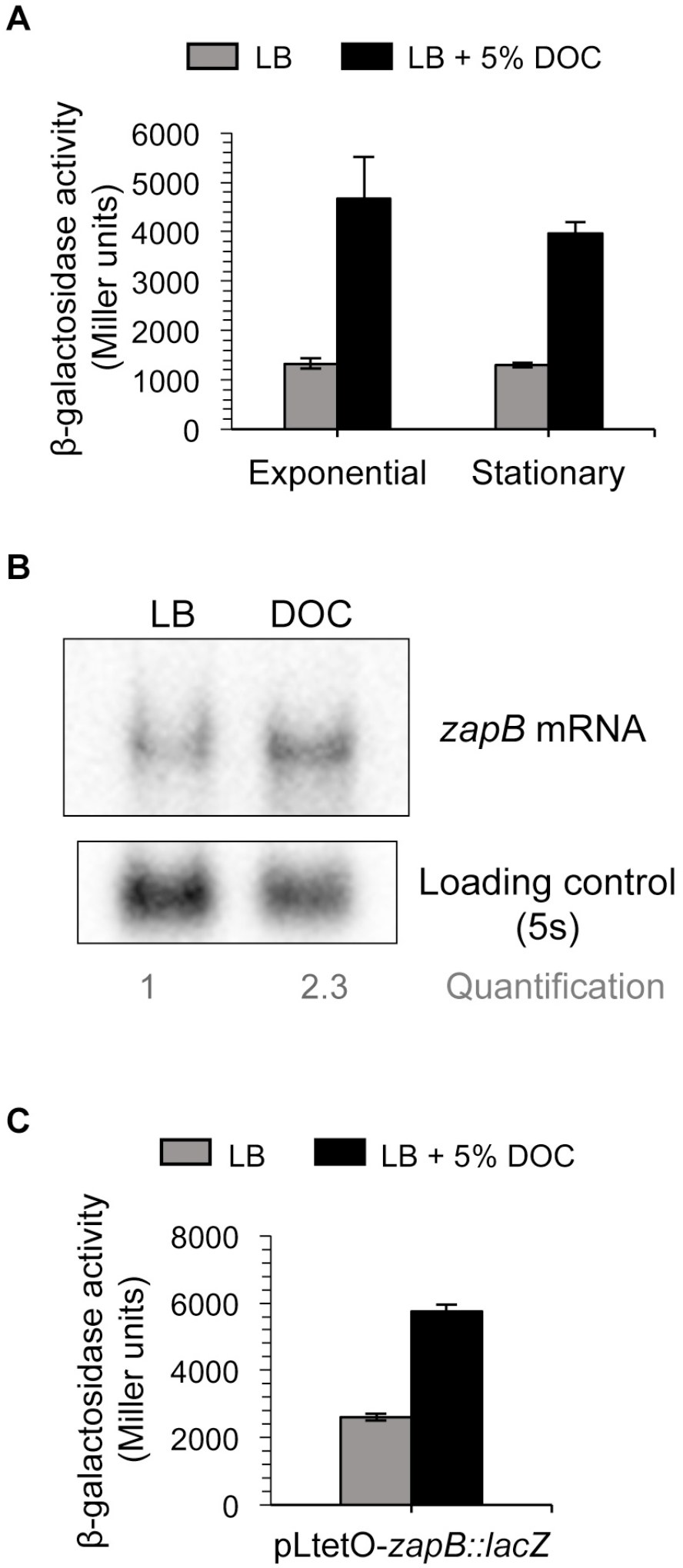
Regulation of *zapB* expression by bile. **A.** β-galactosidase activity of the translational *zapB::lacZ* fusion of strain SV6270 in exponential and stationary cultures in LB (grey histograms) and LB + 5% DOC (black histograms). **B.** Northern blot analysis of the levels of *zapB* mRNA in extracts from stationary cultures in LB and LB + 5% DOC. **C.** β-galactosidase activity of a *zapB::lacZ* fusion under the control of the p_LtetO_ promoter (strain SV7429) in LB (grey histograms) and LB + 5% DOC (black histograms).

To confirm the existence of the predicted *zapB* promoter, the putative promoter region was cloned on the promoter-probe vector pIC552 to generate a transcriptional fusion with the *lacZ* gene. Details of the construction are provided in Suppl. Fig. S1. The pIC552 derivative bearing the *zapB* promoter was introduced into strain SL1344, and β-galactosidase activity measurements confirmed that the cloned region was able to drive *lacZ* expression. However, transcription of p_*zapB*_-*lacZ* was not regulated by DOC (Suppl. Fig. S1). This observation raised the possibility that upregulation of *zapB* expression in the presence of DOC might be postranscriptional. To investigate this possibility, the translational *zapB::lacZ* fusion was placed under the control of an heterologous promoter, pL_tetO_ (32). In this construct, which was engineered on the *S. enterica* chromosome, the native, σ^70^-dependent *zapB* promoter was replaced with pL_tetO_, and the 5’ untranslated region (5’UTR), 56 nt long, was left intact. Measurements of β-galactosidase activity showed that *zapB* expression increased in the presence of DOC from both the native and the heterologous promoter (Fig. 3C), suggesting that DOC-dependent regulation of *zapB* was not transcriptional. This conclusion was in agreement with the above observation that the *zapB* promoter cloned on pIC552 was not responsive to DOC (Suppl. Fig. S1).

### Bile stabilizes *zapB* mRNA by a mechanism involving Hfq

Evidence that upregulation of *zapB* in the presence of bile involved a postranscriptional mechanism prompted a comparison of the stability of the *zapB* transcript in LB and in LB + DOC. For this purpose, Northern blot analysis was performed. We observed that *zapB* mRNA decayed faster in LB than in LB + DOC (Fig. 4A, left pannel), suggesting that the higher level of *zapB* mRNA found in the presence of DOC might result from increased mRNA stability.

**Figure 4.**
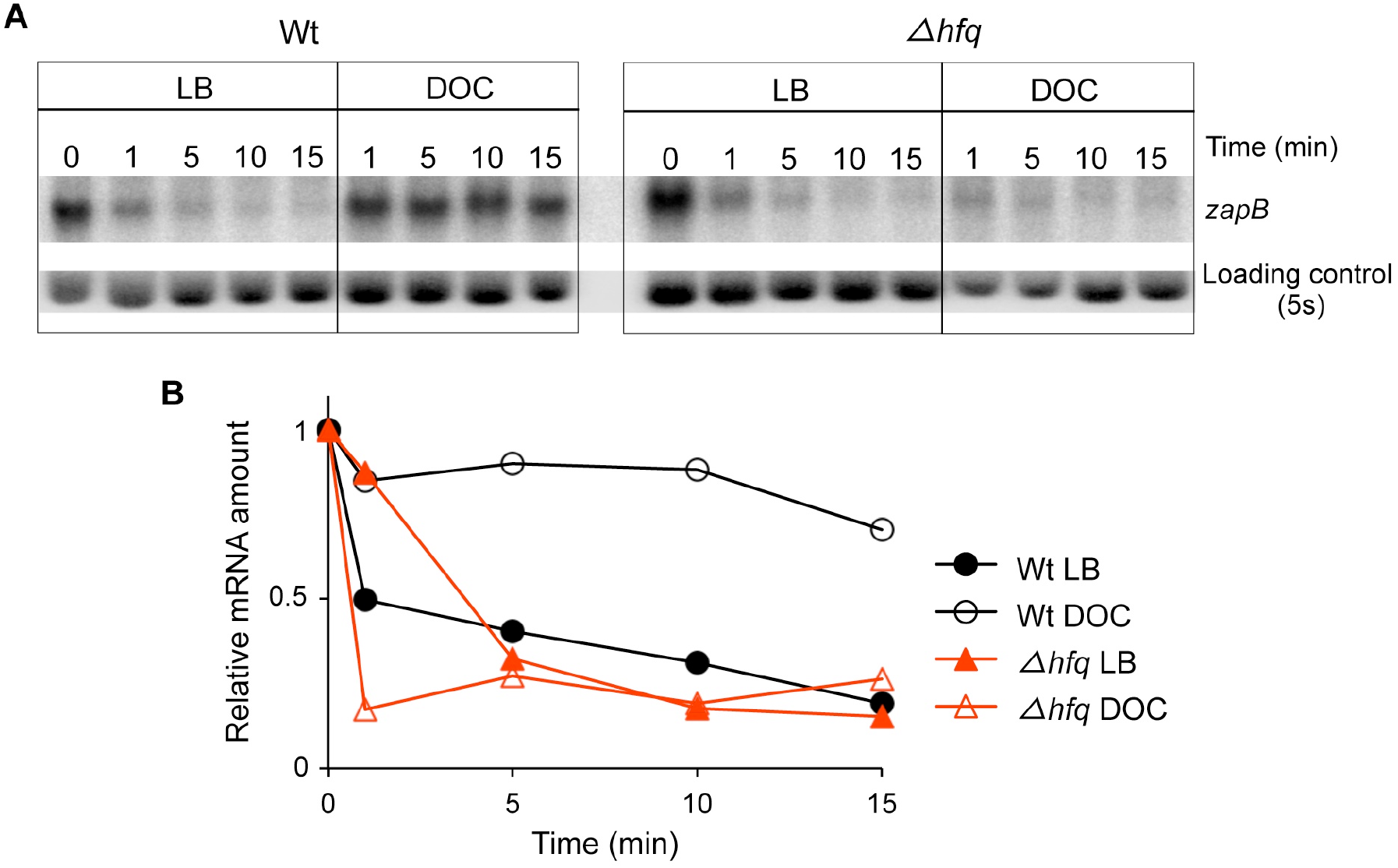
Stability of *zapB* mRNA**. A.** Levels of *zapB* mRNA in RNA extracts from exponential cultures in LB and LB + 5% DOC. Samples were taken 1, 5, 10, and 15 minutes after addition of rifampicin. **B.** Quantification of the level of *zapB* mRNA relative to the loading control and to time zero.

Among the mechanisms that control bacterial mRNA stability, interaction with small regulatory RNAs (sRNAs) is common (33–35). Hence, we considered the possibility that the stability of *zapB* mRNA might be controlled by a sRNA, either destabilizing the transcript in LB or stabilizing the transcript in the presence of DOC. Because many bacterial small regulatory RNAs require the Hfq chaperone for interaction with the target and for stability of the sRNA itself (36–38), we analyzed the stability of *zapB* mRNA in an Hfq^−^ strain (SV5370) using Northern blotting (Fig. 4a, right pannel). In the Hfq^−^ background, the stability of the *zapB* transcript with and without DOC was similar to the stability of *zapB* mRNA in the wild type strain grown in LB, suggesting that stabilization of *zapB* mRNA in the presence of DOC requires the Hfq RNA chaperone (Fig. 4B).

### Identification of the *zapB* mRNA target region involved in postranscriptional control

To identify the *zapB* mRNA region that might be subjected to postranscriptional control, we constructed a set of translational fusions inserting the *lacZ* gene at different locations within the *zapB* coding sequence: 9, 18, 27, 30 and 66 nucleotides after the start codon (Suppl. Fig. S2). Because 3’ untranslated regions (3’UTRs) are often targets for sRNA binding (33, 39), we also constructed a strain in which the 3’UTR of *zapB* was deleted downstream of the *zapB::lacZ* fusion (SV9432). Measurement of β-galactosidase activities upon growth in LB and in LB + DOC provided the following observations:

i. The *zapB::lacZ* fusion that conserved only 9 nucleotides of the *zapB* coding region showed reduced expression both in LB and in LB + DOC (Fig. 5A) and lost regulation by DOC.
ii. DOC-dependent regulation was observed in the remaining *lacZ* fusions (Fig. 5A) including the fusion that maintains 18 nucleotides of the *zapB* coding region, suggesting that the upstream (5’) 18 nucleotides of the *zapB* mRNA coding region are required for regulation by DOC.
iii. The 3’UTR was found to be dispensable for *zapB* regulation by DOC (Fig. 5A). Altogether, the above observations provide evidence that the 5’UTR and the upstream (5’) region of the *zapB* coding sequence may be the main (perhaps the only) region of *zapB* mRNA involved in postranscriptional control by DOC.

**Figure 5.**
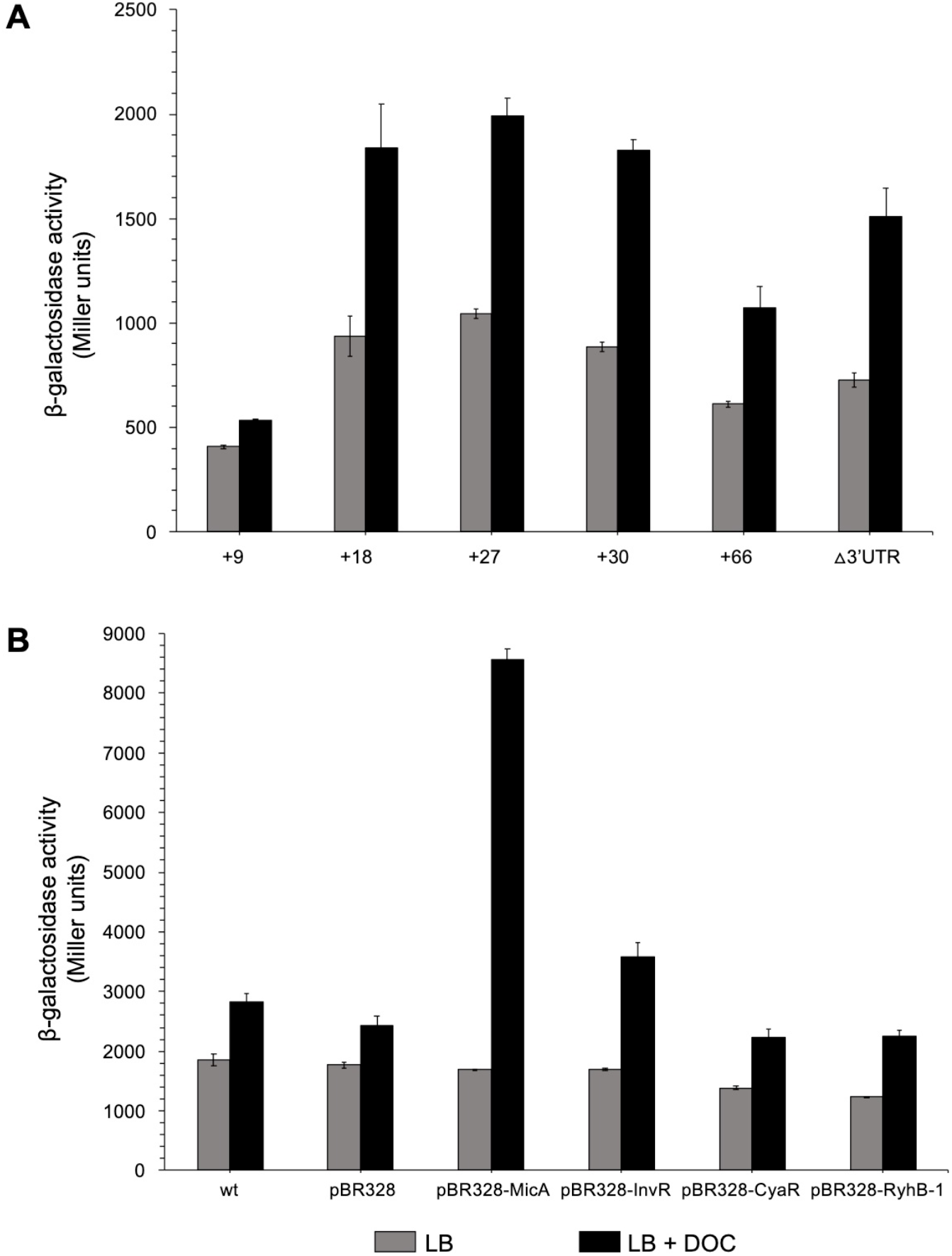
Identification of a *zapB* region involved in regulation by DOC. **A.** β-galactosidase activities of *zapB::lacZ* fusions inserted 9, 18, 27, 30 and 66 nucleotides downstream of the start codon of the *zapB* coding sequence, and of a *zapB::lacZ* fusion lacking the 3’UTR of the *zapB* transcript. **B.** β-galactosidase activities of *zapB::lacZ* fusions in strains overexpressing the sRNAs MicA, InvR, CyaR and RyhB-1 cloned onto the pBR328 plasmid. As controls, the *zapB::lacZ* strain (SV6270, named wild type) and a strain carrying the empty vector were assayed. The strains were grown in LB (grey histograms) and in LB + 5% DOC (black histograms).

### Search for small regulatory RNA(s) that interact with *zapB* mRNA

We did not rule out the possibility that Hfq binding might be sufficient to stabilize *zapB* mRNA. However, this possibility seemed unlikely as Hfq often acts in concert with sRNAs (38, 40–42). A computational search for potential interactions between *Salmonella* sRNAs and *zapB* mRNA using the IntaRNA program (43) identified sRNAs with potential targets in *zapB* mRNA (Supplementary Table S1). Such candidates were shortlisted using two criteria: (i) the potentially interacting mRNA sequence should be located in the 5’ region of the *zapB* transcript; (ii) the potentially interacting sRNA should be regulated by bile. These criteria were met by four candidates: MicA, InvR, CyaR, and RyhB-1, previously reported to be upregulated in the presence of bile (44).

Overexpression systems were set up by cloning MicA, InvR, CyaR and RyhB-1 sRNAs on the multicopy vector pBR328, and transforming the sRNA-carrying plasmids into a *Salmonella* strain harboring a *zapB::lacZ* fusion (SV6270). β-galactosidase activities in LB and in LB + DOC were then determined (Fig. 5B). The presence of DOC resulted in higher β-galactosidase activity when the sRNA MicA was overexpressed. The other sRNAs tested in the assay did not have a significant effect. These results suggest that MicA may be involved in stabilization of *zapB* mRNA in the presence of bile salts.

### Interaction between *zapB* mRNA and MicA

*In silico* analysis with the IntaRNA tool (43) predicts that MicA might interact with *zapB* mRNA by base pairing at a *zapB* mRNA region that contains a putative ribosome binding site (5’GAGG3’) and the AUG start codon of the coding sequence (Fig. 6A). To probe the predicted interaction, point mutations were introduced in MicA sRNA by site-directed mutagenesis. The nucleotide positions and substitutions introduced into the *micA* gene were 5, G→C; 8, G→C; 13, U→A; and 19, A→U. The mutant version of *micA* (*micA*\*) was then cloned on pBR328 and transformed into a strain carrying a *zapB::lacZ* fusion (strain SV9486). Expression of *zapB::lacZ* in LB and in LB + DOC was compared with that of a strain that overexpressed wild type MicA (SV9442). β-galactosidase activity analysis indicated that the upregulation of the *zapB* transcript detected upon MicA overexpression disappeared when MicA was mutated (Fig. 6B). Actually, the β-galactosidase activity in the presence of MicA* was similar to that of the control harbouring an empty vector (Fig. 6B). These observations suggested that the mutations introduced into MicA had disrupted the interaction with *zapB* mRNA, thus supporting the existence of a *zapB* mRNA:MicA sRNA interaction at the region predicted by bioinformatic analysis.

**Figure 6.**
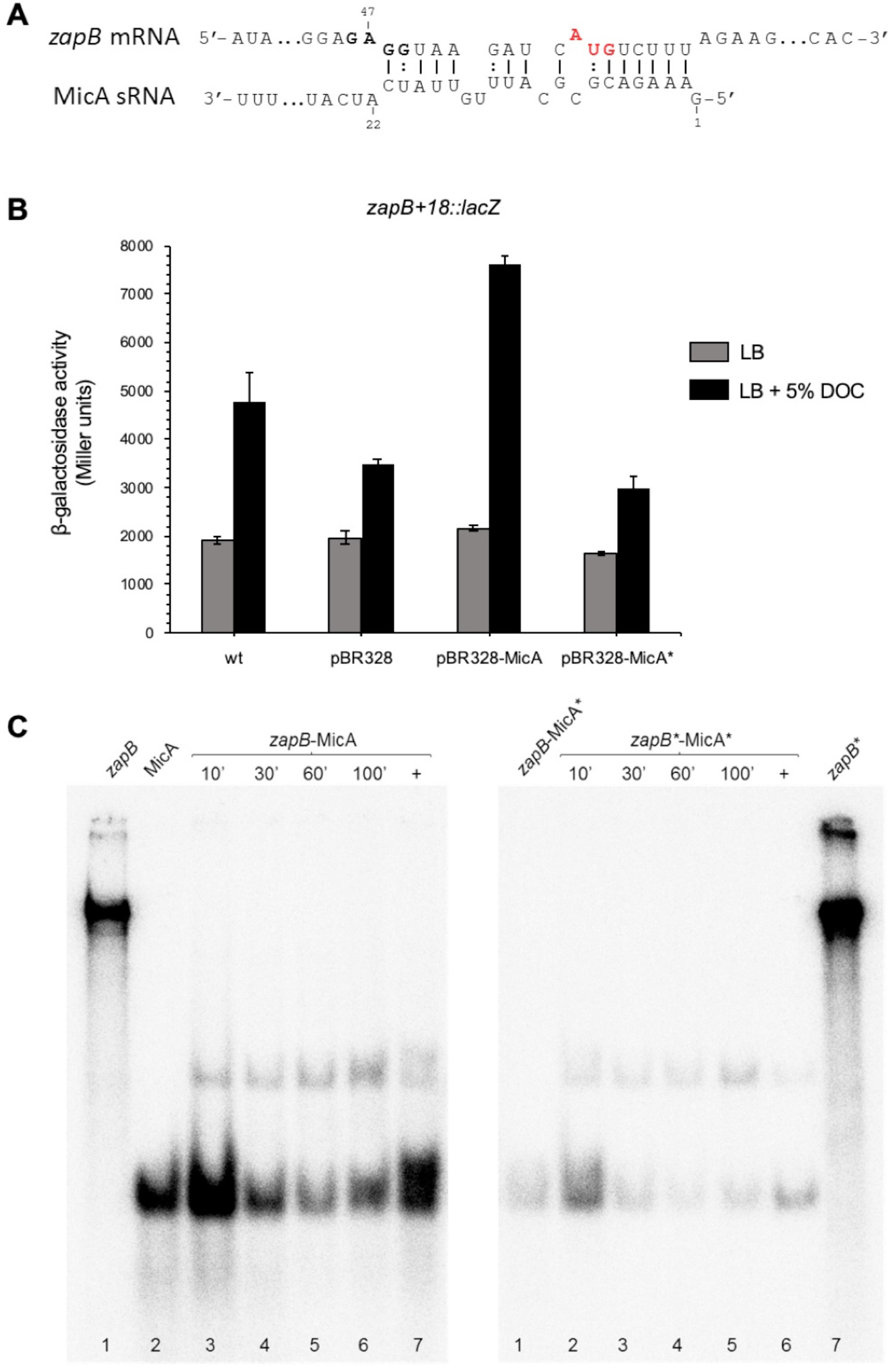
*zapB*-MicA interaction. **A.** Diagram of putative interacting regions in sRNA MicA and *zapB* mRNA predicted by IntaRNA. The putative ribosome binding site and start codon are outlined (bold and red, respectively). **B.** β-galactosidase activities of *zapB::lacZ* fusions in strains overexpressing either MicA or MicA*. As controls, a *zapB::lacZ* strain (SV9372, "wild type") and a strain carrying the empty vector were included. The strains were grown in LB (grey histograms) and in LB + 5% DOC (black histograms). **C.** Gel mobility shift assay (EMSA) showing *in vitro* interactions between *zapB*-MicA, *zapB*-MicA* and *zapB\**-MicA*. Samples were taken at 10, 30, 60 and 100 min. Lanes 1 and 2 in the left pannel and lane 7 in the right panel show labelled *zapB*, MicA and *zapB*\* individual samples, respectively. As positive controls (+), *zapB* and MicA RNAs were treated to force association (lane 7, left and lane 6, right). As negative control, a mix of wild type *zapB* mRNA with mutant MicA (zapB-MicA*) was included (lane 1, right).

Further evidence for *zapB* mRNA:MicA sRNA interaction was obtained *in vitro*. An electrophoretic mobility shift assay (EMSA) was carried out using wild type and mutant versions of both *zapB* mRNA and MicA sRNA, the latter labelled with [α-^32^P] (Fig. 6C). The mutations introduced in *zapB\**, the mutant version of *zapB* mRNA (50: U→A, 54: A→U, 58: U→A, 61, C→G) were complementary to those present in MicA*, so that the interaction predicted by IntaRNA could be restored. Binding was detected for both *zapB* mRNA:MicA and *zapB*\* mRNA:MicA*, and the fact that binding was independent from the incubation time may suggest a strong interaction (Fig. 6C). Altogether, the experiments shown in Figs. 5 and 6 support the following tentative conclusions: (i) that *zapB* mRNA and MicA sRNA do interact; (ii) that the main (perhaps only) interaction involves the upstream region of *zapB* mRNA; and (iii) that *zapB*:MicA interaction increases *zapB* expression in the presence of DOC, probably by MicA-mediated stabilization of *zapB* mRNA.

## DISCUSSION

Initial evidence that the ZapB cell division factor might be involved in bile resistance was provided by transcriptomic analysis: the *zapB* gene, annotated as a locus of unknown function (*yiiU*) in *Salmonella*, was found to be upregulated in the presence of a sublethal concentration of DOC (10). In this study, we show that disruption of the *yiiU* locus causes sensitivity to DOC (Table 1). Hence *yiiU* is not merely a bile-induced locus but a gene necessary for bile resistance. Change of the *yiiU* gene designation to *zapB* is supported by the 91% identity between the predicted *Salmonella* YiiU gene product and the *E. coli* ZapB protein, and by the ability of *E. coli* ZapB to complement sensitivity to DOC in a *S. enterica* YiiU^−^ null mutant (Table 1). Furthermore, the *Salmonella* ZapB protein appears to be localized at the septum like its *E. coli* counterpart (Fig. 1).

Upregulation of *S. enterica zapB* expression in the presence of DOC is still observed when *zapB* transcription is driven by a heterologous promoter (Fig. 3C), thereby suggesting the involvement of a postranscriptional mechanism. Comparison of *zapB* mRNA decay in LB and LB + 5% DOC reveals that *zapB* mRNA is more stable in the presence of DOC, thus explaining the higher mRNA level detected both in the initial transcriptomic analysis (10) and in Northern blots (Fig. 3B). Stabilization of the *zapB* transcript in the presence of DOC requires the Hfq RNA chaperone (Fig. 4), an effect that admits two alternative explanations: (i) Hfq binding might protect *zapB* mRNA from degradation; (ii) Hfq might catalyze the interaction of *zapB* mRNA with a small regulatory RNA, and the mRNA:sRNA interaction might increase *zapB* mRNA stability.

Construction of *lacZ* fusions in truncated forms of the *zapB* transcript defined the 5’UTR and the upstream (5’) 18 nucleotides of the coding sequence as a region necessary for upregulation in the presence of bile salts (Fig. 5A). This hypothesis is in agreement with the fact that activating sRNAs often target upstream mRNA regions (39).

*In silico* search for sRNAs that might interact with *zapB* mRNA through base pairing provided a list of candidates (Suppl. Table S1). Upregulation in the presence of bile (44) and putative interaction at the 5’ region of *zapB* reduced the number of candidate sRNAs down to four: CyaR, InvR, MicA and RyhB-1. When overexpression of these sRNAs was tested, only MicA increased the β-galactosidase activity of a *zapB::lacZ* fusion in the presence of DOC (Fig. 5B). Evidence of a *zapB* mRNA:MicA sRNA interaction was further supported by the existence of a potential base pairing region (Fig. 6A) and by the fact that introduction of point mutations in MicA abolished *zapB::lacZ* upregulation in the presence of DOC (Fig. 6B). Interaction between *zapB* mRNA and MicA sRNA was also detected *in vitro* (Fig. 6C). Site-directed mutagenesis of MicA disrupted the interaction, which was restored upon introduction of compensatory mutations in *zapB* mRNA (Fig. 6C).

MicA is a small regulatory RNA that participates in the bacterial envelope stress response mediated by the sigma factor RpoE (45). MicA binds *omp* mRNAs accelerating their decay and enabling outer membrane remodelling upon envelope stress (46, 47). Identification of additional mRNA targets has linked MicA to the *phoPQ* regulon and to additional functions (48–51). Our observation that MicA protects the *zapB* transcript from degradation in the presence of DOC departs from previous examples of MicA-mediated regulation which involved stimulation of mRNA turnover (46, 47). This difference is further emphasized by the observation that the region of MicA involved in protection of *zapB* mRNA (nt 2-21) overlaps with regions involved in MicA-mediated turnover in other mRNAs: 1-22 in *ompX (50)* and 8-24 in *ompA* (52). These mechanistic differences may seem paradoxical. However, MicA-mediated stabilization of *zapB* mRNA in the presence of bile is anything but paradoxical: a well known role of MicA in the bacterial cell is protection from envelope stress (45–47). Furthermore, transcript stabilization upon interaction with the mRNA 5’ region has been described for other sRNAs (42, 53).

The increased *zapB* mRNA level found in the presence of DOC does not result in higher amounts of ZapB protein (Fig. 2A), and ZapB is degraded by the Lon protease in the presence of DOC (Fig. 2B). Lon degrades abnormally folded proteins (54), and bile salts cause misfolding and denaturation of proteins (6, 7). We thus propose that bile salts may cause ZapB misfolding, and that misfolding may trigger degradation by Lon. If this view is correct, increased *zapB* mRNA stability in the presence of bile may provide a mechanism to compensate for Lon-mediated degradation. Stabilization of *zapB* mRNA may be in turn facilitated by the existence of a higher concentration of MicA in the presence of bile (44).

The cause of bile sensitivity in the absence of ZapB remains unknown. However, it is remarkable that another non-essential cell division factor, DamX, is also necessary for bile resistance in both *S. enterica* (55) and *E. coli* (56). It is thus conceivable that perturbation of Z-ring assembly may render the *Salmonella* cell bile-sensitive.

## MATERIALS AND METHODS

### Bacterial strains, bacteriophages, and culture media

Strains of *Salmonella enterica* serovar Typhimurium (Table S2) derive from SL1344. *Escherichia coli* strains used were RK4353 (*lacUl69 araDJ39 rpsL gyrA*), DH5α [*endA1 hsdR17 supE44 thi1 recA1 gyrA96 relA1* Δ*lacU189* (Ф80 *lacZΔM15*)], CC118 λ *pir* [*phoA20 thi-1 rspE rpoB argE(Am) recA1 (*λ *pir)*] and S17-1 λ *pir* [*recA pro hsdR* RP4-2-Tc::Mu-Km::Tn*7 (*λ *pir)*]. Transduction was performed with *S. enterica* phage P22 HT 105/1 *int201* (57) using a protocol described elsewhere (58). Lysogeny Broth (LB) and M9 were used as liquid media (59). Solid LB contained 1.5% agar. To prepare LB containing sodium deoxycholate (DOC) (Sigma-Aldrich), an appropriate volume from a 25% stock was added. Antibiotics were used at concentrations described previously (60).

### Subcellular fractionation

Separation of cell fractions (cytoplasm, cytoplasmic membrane and outer membrane) was performed as described elsewhere (61).

### Microscopy

Samples grown in M9 were placed on a microscope slide previously spread with 1 µl of poly-L-lysine solution 0.1%. For staining, samples suspended in 100 µl of phosphate buffered saline (PBS) were mixed with 2 µl of Hoechst 33342 (500 µg ml^−1^), incubated 30 min at 30°C and washed with PBS. Images were acquired with a Leica DMR fluorescent microscope and analyzed with Leica IM50 software.

### Immunostaining

Cell preparation and staining were performed as described elsewhere (62). Ethanol-fixed cells (100 µl) were stained with mouse monoclonal anti-FLAG M2-Cy3 antibody (Sigma-Aldrich). Immunostained cells were then stained with Hoechst 33342 in 10 µl mounting medium. Five-ten µl of ethanol-fixed cells were spread onto a poly-L-lysine-coated slide, and dried at room temperature. Slides of stained samples were stored at room temperature in the dark.

### Determination of minimal inhibitory concentration of sodium deoxycholate (DOC)

Aliquots from exponential cultures in LB, each containing around 3×10^2^ colony-forming-units (CFU) were transferred to polypropylene microtiter plates (Soria Genlab) containing known amounts of DOC. After 12 h incubation at 37°C, growth was visually monitored.

### β-galactosidase assays

Levels of β-galactosidase activity were assayed using the CHCl_3_-sodium dodecyl sulfate permeabilization procedure (63).

### Preparation of protein extracts and Western blot analysis

Protein extracts were prepared as described previously (64). For analysis of protein stability, the culture was divided into two: the DOC sample, to which 25% of DOC was added to reach a final concentration of 5%; and the LB sample, to which LB was added to equalize the volume with the DOC sample. A 1 ml aliquot was removed as zero-time control. One ml samples were extracted after 10, 20, 30, 45, 60, 75 and 90 min. Specific proteins were detected with anti-Flag M2 monoclonal antibody (1:5,000) and anti-GroEL polyclonal antibody (1:20,000). Goat anti-mouse horseradish peroxidase-conjugated antibody (1:5,000) or goat anti-rabbit horseradish peroxidase-conjugated antibody (1:20,000) were used as secondary antibodies. Proteins recognized by the antibodies were visualized using luciferin–luminol reagents (Thermo Scientific) in a LAS 3000 Mini Imaging System (Fujifilm). Quantification was performed with MultiGauge software (Fujifilm). GroEL was used as loading control.

### RNA extraction and Northern blot analysis

Aliquots from exponential cultures in LB and LB + 5% DOC were centrifuged at 16,000×g, 37°C, during 2 min, and washed twice with 500 µl of NaCl 0.9%. The pellets were resuspended in 100 µl of a solution of lysozyme (Sigma-Aldrich), 3 mg ml^−1^. Cell lysis was facilitated by freezing at −20°C for >2 hours. After lysis, RNA was extracted using 1 ml of TRIsure reagent (Bioline). Lastly, total RNA was resuspended in 25 µl of RNase-free water. The quality of the preparation and the RNA concentration were determined using a ND-1000 spectrophotometer (NanoDrop Technologies). For Northern blot analysis, 10 µg of total RNA was loaded per well and electrophoresis was performed in denaturing 1% agarose-formaldehyde gels. Vacuum transfer and fixation to Hybond-N^+^ membranes (GE Healthcare) were performed using 0.05 M NaOH. UV crosslinking was used to immobilize RNAs on the membrane. For mRNA stability experiments, rifampicin (500 mg ml^−1^) was added to exponential cultures grown in LB at 37°C (zero-time control). At this point the culture was divided into two: the DOC sample, to which 25% of DOC was added to reach a final concentration of 5%, and the LB sample, to which LB was added to equalize the volume with the DOC sample. Incubation was continued, and culture aliquots were taken at appropriate times. DNA oligonucleotides (Racer2-zapB for *zapB* mRNA and 5S-probe for the loading control) were labelled with [^32^P]-γ-ATP using T4 polynucleotide kinase (Biolabs). As a control of RNA loading and transfer efficiency, the filters were hybridized with an oligoprobe for the 5S rRNA. Membranes were hybridized with oligoprobes at 42°C. Signals were visualized with a FLA-5100 image system (Fujifilm), and quantification was performed using MultiGauge software (Fujifilm).

### 5’rapid amplification of cDNA ends (5’RACE)

RNA was isolated from *S. enterica* stationary cultures (DO_600_ ~2) grown in LB. Fifteen µg of RNA were used to determine cDNA ends using a standard protocol (65). The sequences of the primers are listed in Suppl. Table S3. PCR products were cloned on pGEM-T Easy (Promega), and 29 clones from two independent assays were sequenced.

### Site-directed mutagenesis of *zapB* and *micA*

Introduction of point mutations into MicA sRNA was performed by PCR using as template the wild type *micA* gene cloned on pBR328. Two pairs of primers were employed: MicA5101319-FOR and micAREVSalI; and MicA5101319-REV and micAFORBamHI (Table S3). The resulting DNA fragments were assembled into one using Gibson assembly (66). The fragment was digested with SalI and BamHI and cloned onto pBR328. The plasmid containing substitutions in MicA sRNA (5, G→C; 8, G→C; 13, U→A; and 19, A→U) was transformed into the *zapB*+18*::lacZ* background to yield strain SV9486 (*zapB*+18::lacZ pBR328-MicA*).

To insert point mutations into the chromosomal *zapB* gene, PCR was performed using the pairs of primers For-zapB(MicA)-50545661 and zapB-XbaI-REV; and Rev-zapB(MicA)-50545661 and zapB-SacI-FOR. The two resulting DNA fragments were ligated by Gibson assembly, digested and cloned onto pDMS197 (67). Plasmids derived from pMDS197 were transformed into *E. coli* S17-1 λ *pir*, and the resulting strains were used as donors in matings with the *S. enterica* strain SV9265. Tc^R^ transconjugants were grown in nutrient broth containing 5% sucrose. Individual tetracycline-sensitive segregants were screened for kanamycin sensitivity and examined for the incorporation of the mutant *zapB* allele by DNA sequencing using oligonucleotides zapB-E1 and zapB-E2.

### In vitro transcription and gel mobility shift assay

DNA templates for *in vitro* transcription of *zapB* mRNA and MicA sRNA were generated by PCR using chromosomal DNA from both the wild type and a *zapB* mutant (SV9647), and plasmid DNA from strains containing wild type and mutant variants of sRNA MicA cloned onto pBR328. The primers used were Transc-zapB-FOR and Transc-zapB-REV, and Transc-micA-FOR and Transc-micA-REV (Suppl. Table S3). The PCR products were purified with the Wizard^®^ SV Clean-Up System (Promega). For the *in vitro* transcription reaction, the HiScribe T7 High Yield RNA Synthesis kit was employed. Radioactive labelling was performed by adding 3 µl of EasyTide^®^ uridine 5’-triphosphate (α-^32^P) 3,000 Ci mmol^−1^ (PerkinElmer) to a final concentration of 0.25 µM. Transcription reactions were set up using 1 µl of T7 RNA polymerase per reaction. Nuclease-free water was added to a final volume of 40 µl. The reactions were then incubated at 37°C for 2 h and treated with DNase (TURBO DNase Ambion 2 U µl^−1^) for 15-20 min. To discard unincorporated nucleotides, the reactions were passed through GE Healthcare ilustra™ MicroSpin™ G-25 columns. Subsequently, the RNA transcripts (both labelled and unlabelled) were purified by electrophoresis on a 7.9 M urea, 8% polyacrylamide denaturing gel. The transcripts were visualized using fluor-coated TLC plates (Ambion). Gel slices were crushed, and RNA was eluted overnight at 4°C with crush-soak solution (200 mM NaCl, 10 mM Tris-HCl, 10 mM EDTA). The RNA was ethanol-precipitated and resuspended in nuclease-free water.

Gel mobility shift assays were performed with 0.015 pmol of [α-^32^P]-labelled wild type or mutant MicA (MicA*) RNA, and 0.075 pmol of unlabelled *zapB* mRNA transcript in 1× binding buffer (200 mM Tris-HCl pH 8, 10 mM DTT, 10 mM MgCl_2_, 200 mM KCl, 100 mM Na_2_HPO_4_-NaH_2_PO_4_) at 37°C. The reactions were incubated, and samples were taken at 10, 30, 60 and 100 min. The binding reactions were mixed with 2 µl of loading dye (48% glycerol, 0.01% bromophenol blue) and subjected to electrophoresis at room temperature in a 6% nondenaturing polyacrylamide Tris-borate EDTA (TBE) gel for 2 h at 55 V, 15 mA in 1x TBE buffer (90 mM Tris-borate/2 mM EDTA). In parallel to the binding reactions, a positive control was prepared by mixing *zapB* mRNA and [α-^32^P]-labelled MicA (mutant and wild type) followed by boiling denaturation and renaturation by incubation at 37°C. Additionally, single [α-^32^P]-labelled *zapB* RNA and MicA were loaded on the gel. After electrophoresis, gels were dried and analyzed in a FLA-5100 Scanner (Fujifilm).

## ACKNOWLEDGMENTS

This study was supported by grant BIO2016-75235-P from the Spanish Ministerio de Economía y Competitividad (MINECO). V. U. was supported by a postdoctoral contract associated to grant CVI-5879 from the Consejería de Innovación, Ciencia y Empresa, Junta de Andalucía, Spain. We are grateful to Miguel A. de Pedro and Kai Papenfort for discussions, and to Modesto Carballo, Laura Navarro, and Cristina Reyes of the Servicio de Biología, CITIUS, Universidad de Sevilla, for help with experiments performed at the facility.

